# Quantitative 2D J-resolved metabolite-cycled semiLASER spectroscopy of metabolites and macromolecules in the human brain at 9.4 T

**DOI:** 10.1101/2024.09.30.615877

**Authors:** Saipavitra Murali-Manohar, Tamas Borbath, Andrew Martin Wright, Nikolai I. Avdievich, Anke Henning

## Abstract

**Purpose:** While two-dimensional (2D) in vivo spectroscopy yields rich information and has been successfully used in clinical trials, it requires a localization scheme that minimizes the impact of chemical shift displacement on J-coupling evolution, a robust frequency drift correction and dedicated processing and quantification methods. Considering these needs this study demonstrates a novel data acquisition and an analysis pipeline to quantify 16 metabolites in mmol/kg in the human brain using a 2D J-resolved metabolite-cycled (MC) semiLASER localization sequence at 9.4 T in the human brain.

**Methods:** Metabolite spectra were acquired in vivo using the newly developed J-resolved MC semiLASER localization sequence with maximum echo sampling (MES) at 9.4 T. In order to account for the underlying macromolecular (MM) spectra in the acquired metabolite spectra, J-resolved MM spectra were acquired using a double inversion recovery (DIR) J-resolved MC semiLASER. Spectral fitting was performed with ProFit 2.0 using a simulated basis set from VesPA tailored to 2D J-resolved semiLASER with MES. Finally, metabolite concentrations were calculated using internal water referencing.

**Results:** Tissue concentrations for 16 metabolites in mmol/kg are reported after correcting for number of protons, tissue content, and relaxation effects of both water and metabolites at 9.4T. Quantification results of spectra considering 8 and 2 averages per TE did not show any significant differences.

**Conclusion:** 2D spectra of metabolites acquired at 9.4T and 2D MMs acquired at any field strength are presented for the first time. Basis set simulation and quantification of metabolites for metabolite spectra acquired using maximum-echo-sampled 2D J-resolved semiLASER was performed for the first time. The sensitivity in the detection of J-coupled metabolites such as glutamine, glucose or lactate. At ultra-high field, the acquisition duration of 2D MRS can be also substantially reduced since only a very low number of averages per TE are needed.

## 1. Introduction

Single voxel proton magnetic resonance spectroscopy (^1^H-MRS) is a popular non-invasive technique used to study the metabolism in the human brain. It enables detection and quantification of the neurochemical profile in a particular region of interest, thereby aiding detection of various pathologies in the human brain^1^. However, complex spectral patterns and severe spectral overlap often pose a challenge in quantifying individual metabolite concentrations^2^. Therefore, enhancing the signal-to-noise ratio (SNR) and spectral resolution has always been one of the main aims of the MRS community.

One technique used by the NMR community to reduce spectral overlap is multi-dimensional spectroscopy^3,4^. Homonuclear two-dimensional (2D) techniques such as correlation spectroscopy and J-resolved spectroscopy were shown to be feasible in vivo at 1.5, 3 and 7 T^5-7^ and hold the promise to yield accurate quantification results for a large number of metabolites in clinical trials of neurological and psychiatric disorders^8-14^. Nevertheless, at clinical field strength this technique owes to longer measurement durations than 1D MRS with 4 to 8 averages needed for each encoding step and requires non-standard pre-processing, fitting and quantification routines.

In 2D J-resolved spectroscopy^15^, the spectral information is spread along two orthogonal axes by adding a step-wise increasing evolution delay in the pulse sequence. This helps improve specificity in the detection of J-coupled metabolites. To further improve detection sensitivity, maximum echo sampling (MES)^16^ was introduced for in vivo 2D J-resolved ^1^H MRS^8,17^. In the MES scheme, acquisition begins right after the final crusher gradient of the last refocusing pulse in the localization technique. Schulte et al^15^ showed that JPRESS with MES had increased sensitivity in comparison to traditional half echo sampling. Yet another advantage of MES is that it adds a tilt to the peak tails in the spectrum^15^ and thus largely reduces overlap of the water peak tail with the metabolite peaks of interest.

Another means to clearly distinguish more peaks is ultra-high field (UHF) (≥ 7 T) ^1^H MRS^18,19^. It is well known that spectroscopy studies at UHF benefit from both increased spectral resolution and improved SNR in comparison to lower field strengths. However, at UHF B_1_^+^ inhomogeneity is one of the crucial challenges posed. Also, with increasing static B_0_ field strength, chemical shift displacement error between the metabolite peaks increases. Furthermore, quantification for J-coupled peaks such as glucose is easier at a lower field strength as shown previously^20^.

Using simulation and experimental methods at 3 T and 7 T Edden et al^21^ demonstrated that the chemical shift displacement effect causes spatially dependent differences in J-evolution of coupled spin systems in JPRESS spectra. Due to the larger spectral dispersion along with reduced peak transmit field strength B_1_^+^, the chemical shift displacement error is increasing with increasing field strength. The appearance of additional J-refocused peaks results in loss of intensity in the J-resolved peaks and more spectral overlap thereby leading to uncertain spectral quantification. However, Lin et al^6^ compared the spectral quality of JPRESS, J-resolved semiLASER and J-resolved LASER sequences at 3 T and demonstrated that the use of adiabatic pulses reduced the appearance of J-refocused peaks to a great extent due to their higher bandwidth. Adiabatic pulses in the localization schemes are also effective in reducing the effect of B_1_^+^ inhomogeneity at UHF^22^. Both the implications discussed above emphasize the need to use adiabatic localization when implementing J-resolved spectroscopy at UHF.

Finally, the metabolite cycling (MC) scheme^23^ enables simultaneous acquisition of water and metabolite signals. It has been shown for 1D MRS at 3T^24-26^, 7T^27^ and 9.4T^28^ previously that retrospective frequency alignment based on the MC water signal substantially decreases the linewidth of the resulting spectra yielding a complimentary mechanism to enhance the spectral resolution and data consistency.

Therefore, in this work, we developed a 2D J-resolved metabolite-cycled (MC) semiLASER sequence for human brain application at 9.4 T. Implementing MC^23,28^ for J-resolved spectroscopy eliminates the need for acquiring a water reference scan for the sake of eddy current correction or frequency alignment which leads to reduction of scan time. An added bonus is that one can also simultaneously measure both upfield and downfield part of the ^1^H spectra when employing MC^28,29^.

Even though, the 2D J-resolved semiLASER acquisition method was developed earlier at 3 T^6^, metabolite quantification was not performed. This work includes basis set simulation for 2D J-resolved semiLASER for the first time. In addition, spectral fitting of 2D spectra (considering 8 and 2 averages per TE) was performed using ProFit v2.0^30^, a dedicated 2D fitting software, which incorporates the theory of LCModel^31^ and VARPRO^32^ in attaining a global minimum. Fuchs et al^23^ enhanced the software by adding a macromolecular model, a spline baseline fit and a spline lineshape model using self-deconvolution. Spectra were considered with 8 and 2 averages per TE for processing, fitting and quantification in order to evaluate if there is significant difference in the concentration values between the low field standard and the accelerated acquisition scheme at UHF.

Subsequently, quantification of metabolites was performed using the internal water referencing method^33^ taking into account the tissue composition in the voxel of interest, and water concentration. Finally, quantification results obtained with 2D J-resolved spectroscopy were compared against metabolite concentrations published in previous 1D MRS studies. To the best of our knowledge, the effect of bringing together the two complementary approaches of UHF and 2D J-resolved spectroscopy to enhance spectral resolution on spectral quantification has not been investigated yet. This work is an extension of the initial results reported earlier in a conference abstract^34^.

## 2. Methods

### 2.1 Study Design

All measurements were performed on a 9.4 T Siemens Magnetom whole-body MRI scanner (Siemens Healthineers, Erlangen, Germany) using a home-built coil^35^ with eight transmit and sixteen receive channels. Eleven healthy volunteers (6 male and 5 female, age: 28.0 ± 2.3 years) participated in this study. Five volunteers returned for a second visit for the acquisition of 2D MM signal. The study was approved by the local ethics board, and written informed consent was provided by the volunteers prior to the measurements.

### 2.2 MRS Sequence

Figure 1a shows the J-resolved metabolite-cycled (MC) semiLASER sequence diagram. The asymmetric adiabatic MC^28^ pulse (duration: 22.4 ms) preceded the conventional semiLASER^28,36^ block. A hamming filtered 90 degree-sinc pulse^28^ (bandwidth: 8000 Hz) was used for excitation. This pulse was followed by two pairs of adiabatic full passage pulses (duration: 3.5 ms, bandwidth: 8000 Hz). The indirect dimension (t_1_) was created by inserting an incrementally increasing time delay of Δt/2 between the last pair of adiabatic full passage pulses^28^, which encodes the J-evolution. The acquisition of the signal began immediately after the final crusher gradient of the last adiabatic full passage pulse which is called maximum echo sampling (MES) scheme^15^.

**Figure 1.**
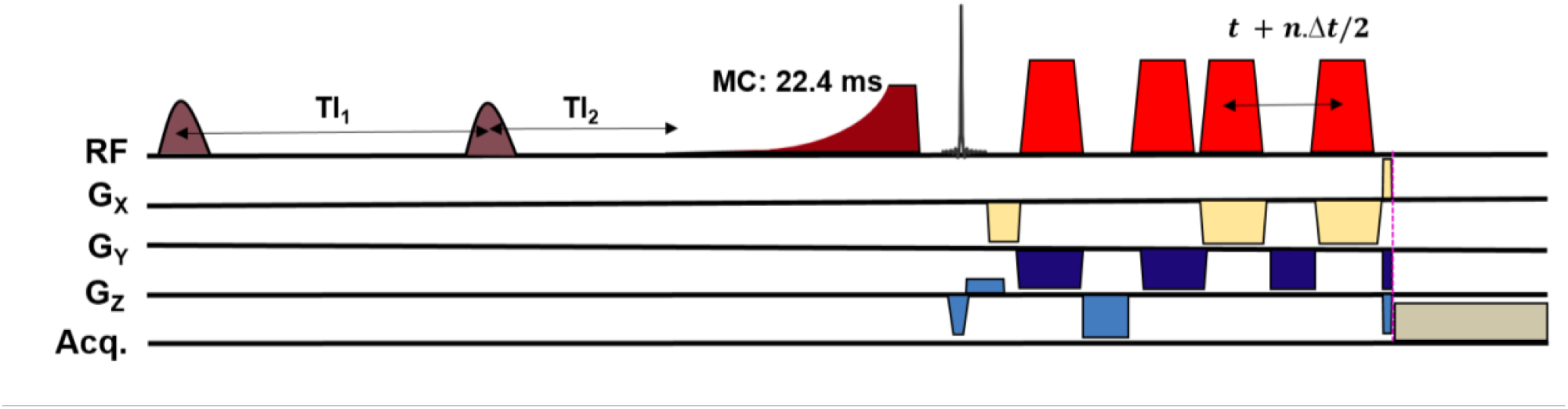
Pulse sequence diagram of J-resolved metabolite-cycled semiLASER localization scheme. The two inversion pulses were turned off when acquiring the metabolite spectra. For the acquisition of macromolecular spectra, double inversion recovery technique was used with TI_1_/TI_2_ set to 2360/625 ms. The indirect dimension is created by increasing the time interval between the last two AFP pulses by Δt/2. Both the sequences have maximum echo sampling scheme implemented i.e., acquisition begins right after the final crusher gradient of the last AFP.

In order to optimize the number of TE steps *n*, knowledge of T_2_ relaxation times of metabolites in vivo at 9.4 T was utilized from Murali-Manohar et al^37^ (further explained in section 4.1). Additionally, phantom measurements (Figure 2) were performed with different time increment steps (Δt: 2, 3, and 4 ms) and different number of TE steps (*n*: 50, 85) to confirm the absence of J-refocused peaks and whether the peaks are distinctly J-resolved without noise and truncation artifacts in the indirect dimension.

**Figure 2.**
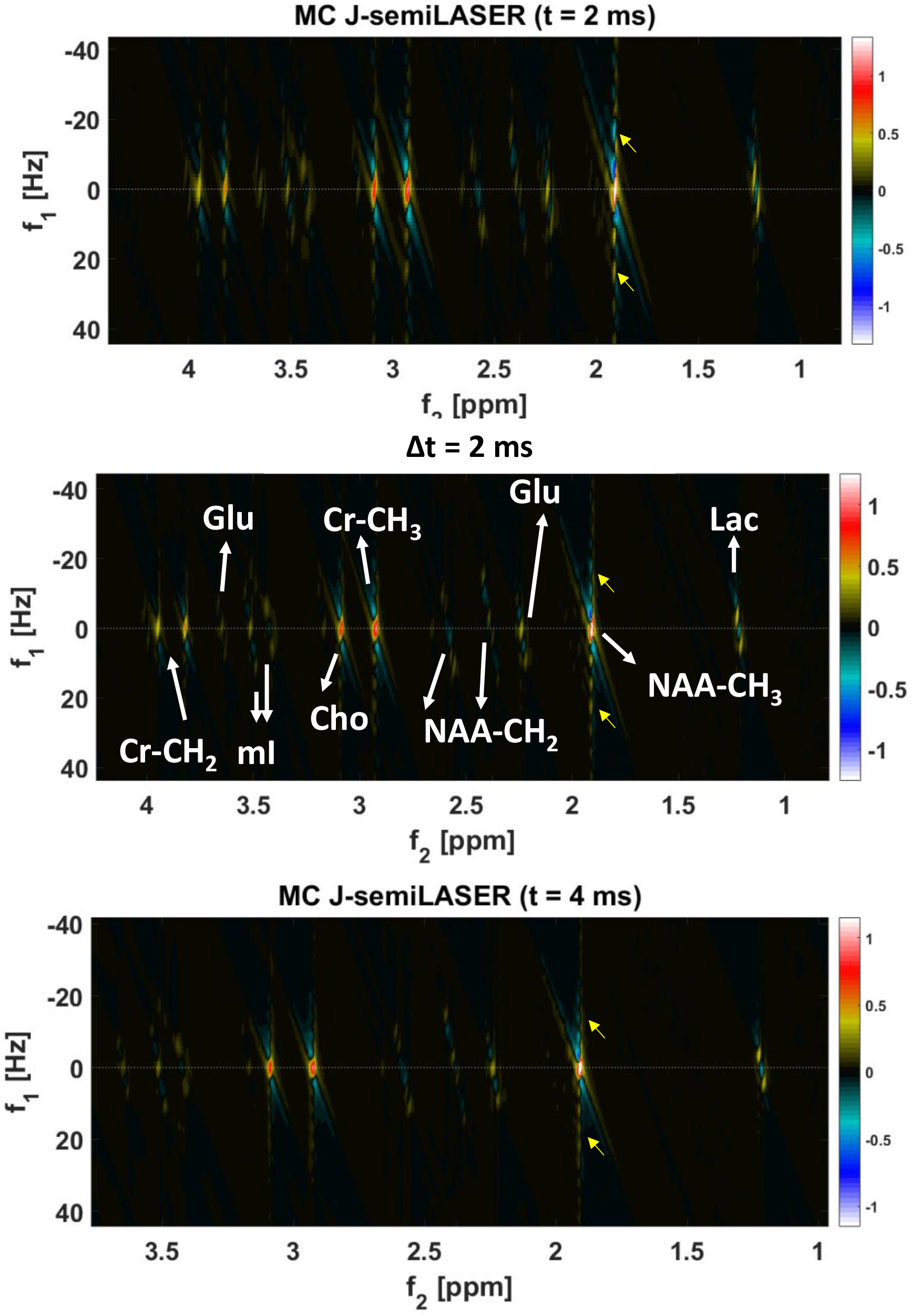
(Top to bottom) 2D J-resolved MC semiLASER spectra from Braino phantom containing NAA, Cr, Cho, Glu and Lac with n = 50 and Δt = 2, 3 and 4 ms. White arrows indicate the t_1_-ridges as a result of t_1_ noise and truncation sinc artifacts in the indirect dimension. The ridges are strongly pronounced when TE_max_ = 124 ms and minimum when TE_max_ = 224 ms.

A double inversion recovery (DIR) block preceding the J-resolved MC semiLASER sequence (Figure 1b) was used for the acquisition of macromolecular signal (*n*: 60). The inversion pulse^38^ duration was 15 ms and the inversion bandwidth was approximately 1650 Hz.

### 2.3 Data Acquisition

#### 2.3.1 Anatomical Imaging

MP2RAGE^39^ images (resolution: 0.6 × 0.6 × 0.6 mm^3^) were acquired while using the coil in volume mode driving power to all the eight transmit coil elements.

For the MM acquisition, only 2D FLASH images were acquired which then were co-registered to the previously acquired MP2RAGE images using rigid body transformation.

#### 2.3.2 Spectroscopy Measurements

Following the anatomical scan, the volunteer was instructed to remain stationary on the patient table while the coil setup was changed to suit the spectroscopy measurements. Power was driven to the bottom three coil elements near the region of interest using unbalanced three-way Wilkinson splitter^28^. A localizer was reacquired to ensure there was no motion of the volunteer between the anatomical and the spectroscopy scan. This was followed by acquisition of high-resolution 2D FLASH images (in-plane resolution: 0.7 × 0.7 mm^2^, slice thickness: 3.5 mm, 25 slices) in the sagittal and transversal orientations to position the spectroscopy voxel (2 × 2 × 2 cm^3^) in the occipital lobe. Localized second-order shimming was performed using FAST(EST)MAP^40^ setting the shim volume to be 150% of the volume of the voxel of interest. Then, voxel-based power optimization^41,42^ was performed on each subject to ensure that the adiabatic conditions were fulfilled.

Two-dimensional metabolite spectra were acquired using the J-resolved MC semiLASER sequence (Figure 1) (TR: 6000 ms, averages per TE step: 8, four-step phase cycling, transmit reference frequency: 2.4 ppm) described above in the MRS sequence section. The MC pulse had a duration of 22.4 ms, frequency factor 2 and offset frequency ± 350 Hz^28^. TE ranged from 24 to 194 ms (*n*: 85) incremented in steps of Δ*t* = 2 ms. In addition, 2D water reference signals were acquired (average per TE step: 1, transmit reference frequency: 4.7 ppm) for validation purpose to avoid any saturation effect of the water due to the MC pulse for the quantitative evaluation. The transmit reference frequency for the water reference sequence was set to 4.7 ppm.

In order to account for the MM contribution in the 2D metabolite spectra, 2D MM spectra were acquired from five healthy volunteers using the DIR sequence. The optimized inversion times^30,35^ (TI_1_/TI_2_) combination of 2360 and 625 ms was used in the DIR block as shown in Figure 1b. TE ranged from 24 to 144 ms (*n*: 60) since MM signal decays faster in comparison to metabolite signal^29^ and 16 averages were acquired at each TE step. All the other acquisition parameters were identical to the metabolite spectra except the TR which was set to 8000 ms^43^ to fulfill SAR constraints.

### 2.4 Data Preprocessing

Spectroscopy raw data were preprocessed using an in-house MATLAB (version 2016a, MathWorks, Natick, MA) tool. Both metabolite and MM data were reconstructed as described in Giapitzakis et al^28^. Firstly, the data were frequency and phase-aligned based on the water signal in the time domain for 85 and 60 blocks for metabolite and MM data respectively. This was followed by metabolite-cycling subtraction and then the data were averaged within each TE block considering 8 and 2 averages per TE. Then Eddy current correction was performed using the phase information from the MC water signal. Signals from all 16 receive channels were then combined using the SVD method. The metabolite and MM data were truncated at 250 and 150 ms respectively for better SNR and considering their T_2_ relaxation times at 9.4 T. Automated zeroth- and first-order phase corrections were performed in the J-resolved spectroscopy preprocessing tool^15,36^ which is a part of ProFit. The applied phase correction was visually verified for correctness as recommended by the recent consensus article^37^. Using the same ProFit preprocessing tool, residual water in both the dimensions in the spectra was removed using a HSVD method. The final 2D spectrum displayed is after applying Fourier transformation to the data in both the dimensions. SNR of the NAA(CH_3_) peak at 2.008 ppm was calculated as the peak intensity from the real part of the spectrum with respect to the noise window from -4.0 to -1.0 ppm.

### 2.5 MP2RAGE Segmentation

The high-resolution MP2RAGE images were segmented into gray matter (GM), white matter (WM) and cerebrospinal fluid (CSF) fractions using SPM12^38^. 2D FLASH images acquired during the MM scan session were co-registered to the MP2RAGE images acquired during the first scan session for four volunteers. A home-built Python (v3.7) tool was used to determine the tissue fractions in the voxels of interest.

### 2.6 Spectral Fitting

Metabolite basis sets corresponding to 85 TE steps were simulated using VesPA (ver. 0.9.3)^44^ considering full quantum mechanical density matrix calculations for the semiLASER sequence including the excitation and adiabatic full passage pulse shapes. The simulation also incorporated the sequence timings including the MES scheme. The following metabolites were simulated: n-acetyl aspartate (NAA), NAA glutamate (NAAG), *γ*-aminobutyric acid (GABA), aspartate (Asp), creatine (Cr), glutamate (Glu), glutamine (Gln), glucose (Glc), glutathione (GSH), glycerophosphocholine (GPC), glycine (Glyc), myo-inositol (mI), scyllo-inositol (Scy), lactate (Lac), phosphocreatine (PCr), phosphocholine (PCho), phosphoethanolamine (PE), and taurine (Tau). Their chemical shifts and coupling constants were chosen according to Govindaraju et al^39^, except for the coupling constant of GABA, for which the values from Near et al^40^ were chosen. Subsequently different metabolite peaks were scaled to correct for the number of proton contributions. GPC, PCho and PE combined together is denoted by tCho. Finally, all the 85 1D basis sets and the measured MM spectra were combined to form a complete 2D basis set using an in-house written MATLAB script.

All the metabolite spectra (8 and 2 averages per TE) were fitted using ProFit 2.0^41^ (2D PRiOr knowledge FITting). It takes the exponential decay of the metabolite signal and the scalar coupling constants into account and fits the J-resolved spectrum after 2D Fourier transform in both the direct and the indirect dimensions. The non-linear fitting routine iterated four times, each time with increasing degrees of freedom.

### 2.7 Quantification

The formula used for the quantification of metabolites^33^ is provided in the Supplementary Material Annex A. Wilcoxon signed rank tests were conducted for each metabolite measurement with 8 averages and 2 averages per TE (α/m = 0.003846) and adjusted-p values were calculated.

## 3. Results

### 3.1 Voxel Content and Spectral Quality

The spectra from the phantom tests (*n*: 50, Δ*t*: 2, 3, and 4 ms) showed stronger ‘t_1_ ridges’^21^ when TE_max_ was shorter (Figure 2). In other words, when TE_max_ = 124 ms, the ‘t_1_ ridges’^21^ were the strongest. t_1_ noise was the least when TE_max_ = 224 ms. When TE_max_ was set to 100, 99, and 100 ms for Δ*t*: 2 ms (*n*: 50), 3 ms (*n*: 33), and 4 ms (*n*: 25) respectively, the ‘t_1_ ridges’ were not seen to be impacted (Supplementary figure S1) when changing the delay between TE steps alone. However, Δ*t*: 2 ms corresponded to higher SNR since it allows for a greater number of averages for a chosen TE_max_ and was hence used for all additional experiments since a good SNR is essential for in vivo spectra. Even though the t_1_ noise seemed to be minimum for a TE_max_ = 224 ms, TE_max_ = 194 ms was chosen for in vivo studies as T_2_ relaxation times are generally longer in phantom than in in vivo tissues.

Figure 3 shows a phantom metabolite spectrum (*n*: 85, Δ*t*: 2 ms, TE_max_: 194 ms) in the magnitude mode. There are neither prominent J-refocused peaks appearing in the spectrum nor are there any visible truncation artifacts or t_1_ noise in the indirect dimension.

**Figure 3.**
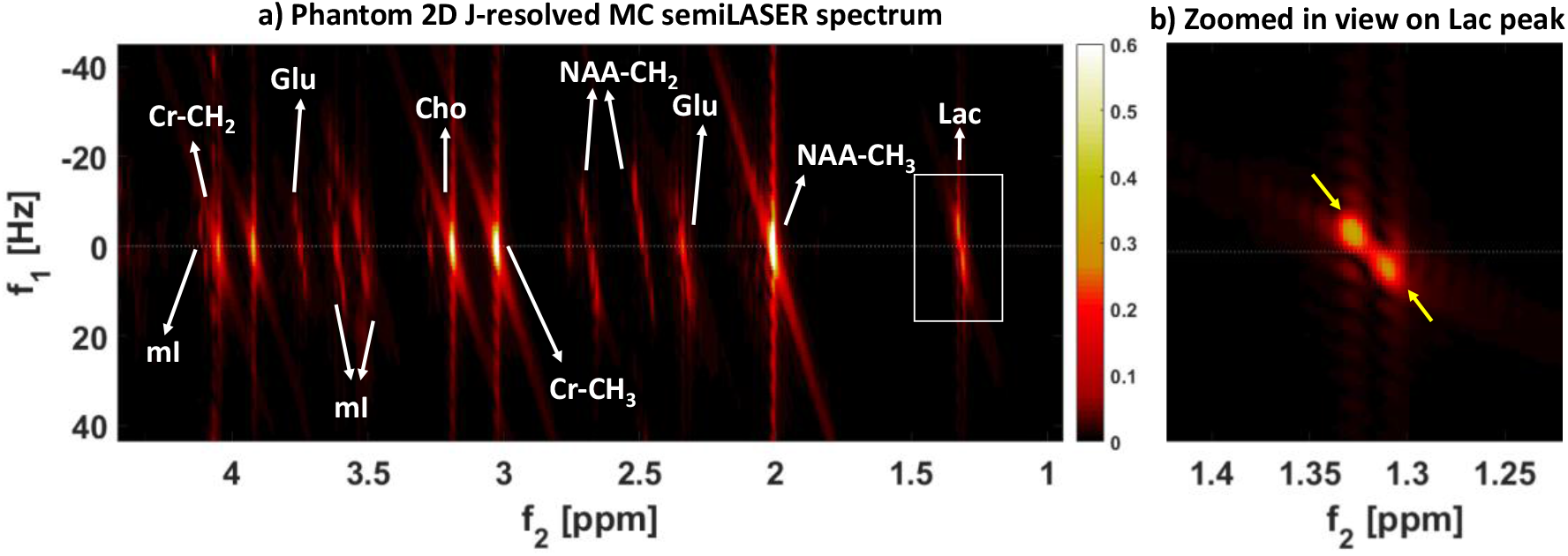
a) 2D metabolite spectrum from Braino phantom (*n* = 85, Δ*t* = 2ms) in magnitude mode b) Zoomed in view of the lactate peak showing a J-resolved doublet but barely any J-refocused peaks. An 86% suppression (considering Lac peaks at 1.31 and 4.09 ppm since they have a maximum separation of values in the observable range of the spectrum) of the J-refocused peaks is expected when calculated as suggested by Lin et al^6^ resulting in barely any visible J-refocused peaks. This in turn leads to maximum intensity of the J-resolved peaks.

Figure 4 shows a representative single subject 2D MC J-resolved semiLASER metabolite spectrum (*n*: 85, Δ*t*: 2 ms, 8 averages per TE) with an inlay showing the voxel position. Figure 7 shows the respective 2D J-resolved MC semiLASER spectrum considering only 2 averages per TE. The average tissue content of the metabolite spectroscopy measurement voxel in the occipital lobe of eleven healthy volunteers was GM/WM/CSF = 67 ± 8/ 29 ± 9/ 4 ± 1 % respectively. The 2D metabolite spectra obtained from voxels in the occipital lobe showed uncoupled peaks lying on the f_1_ = 0 Hz axis. This is because during t_1_, only the coupling information is obtained. However, the J-coupled multiplets are resolved at an angle of 45° with respect to f_1_ = 0 Hz since t_2_ holds both chemical shift and coupling information. The SNR of the NAA(CH_3_) peak in case of 8 averages per TE step was 906 ± 147 and in case of 2 averages per TE step was 648 ± 105 indicating that both the 2-average and the 8-average spectra were of good quality. Also, the spectra did not show any major artifacts such as lipid contamination or water tails. Therefore, no data sets were excluded from further analysis.

**Figure 4.**
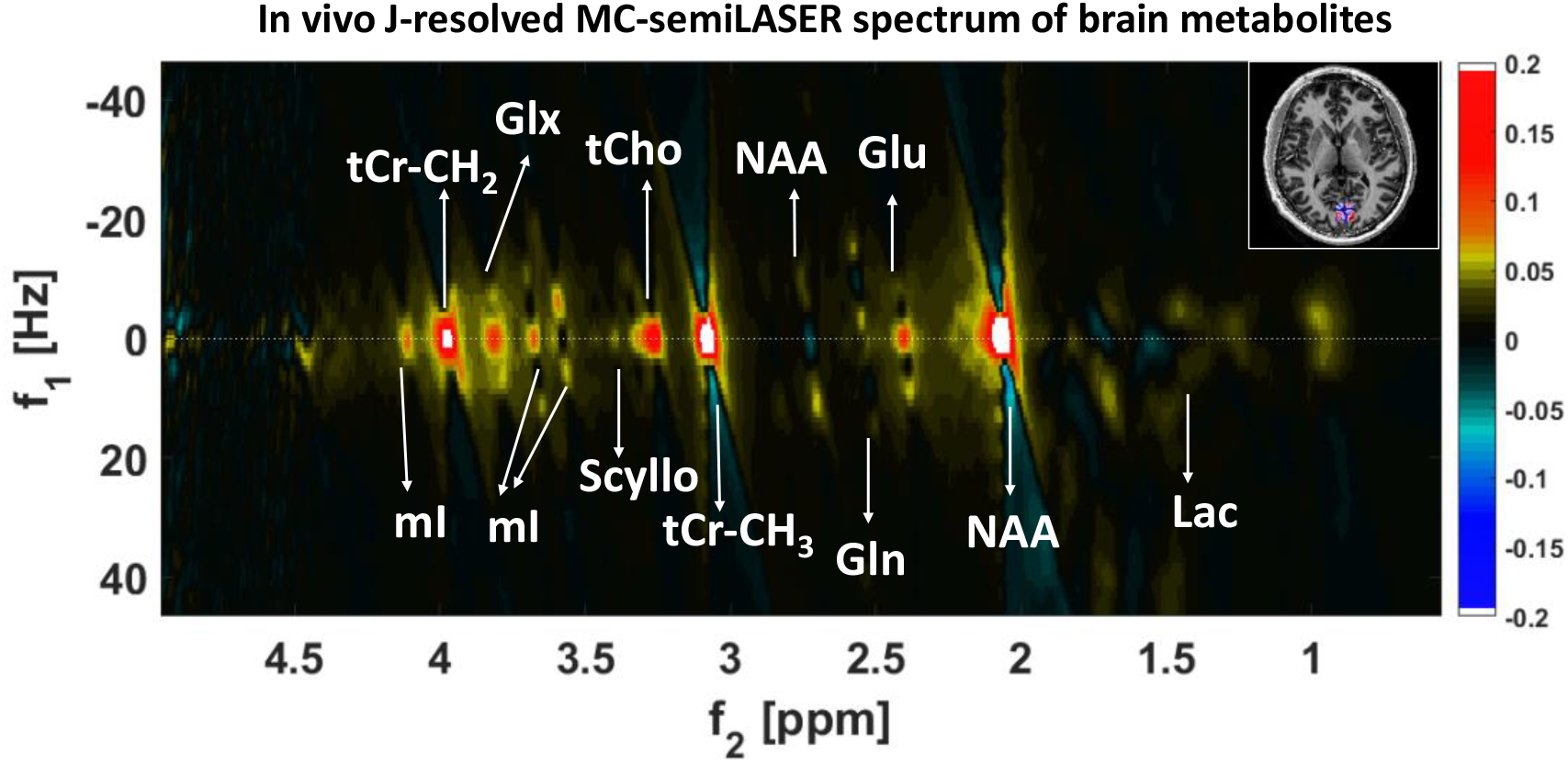
2D J-resolved MC semiLASER spectrum from a representative subject with *n* = 85 and Δ*t* = 2 ms. The figure inlay shows the voxel positioning on the MP2RAGE image in transversal view. Additionally, the observed metabolites are labeled in the figure.

The summed MM spectrum from five healthy volunteers is shown in Figure 5. M_0.92_, M_1.21_, M_1.39_, M_1.67_, M_2.04_, M_2.26_, M_2.56_, M_2.70_, M_2.99_ and M_3.86_ are clearly seen to have J-resolved peaks. On an average, the MM spectroscopy voxel (five healthy volunteers) had GM/WM/CSF = 61 ± 10/ 35 ± 8/ 4 ± 3 % respectively.

**Figure 5.**
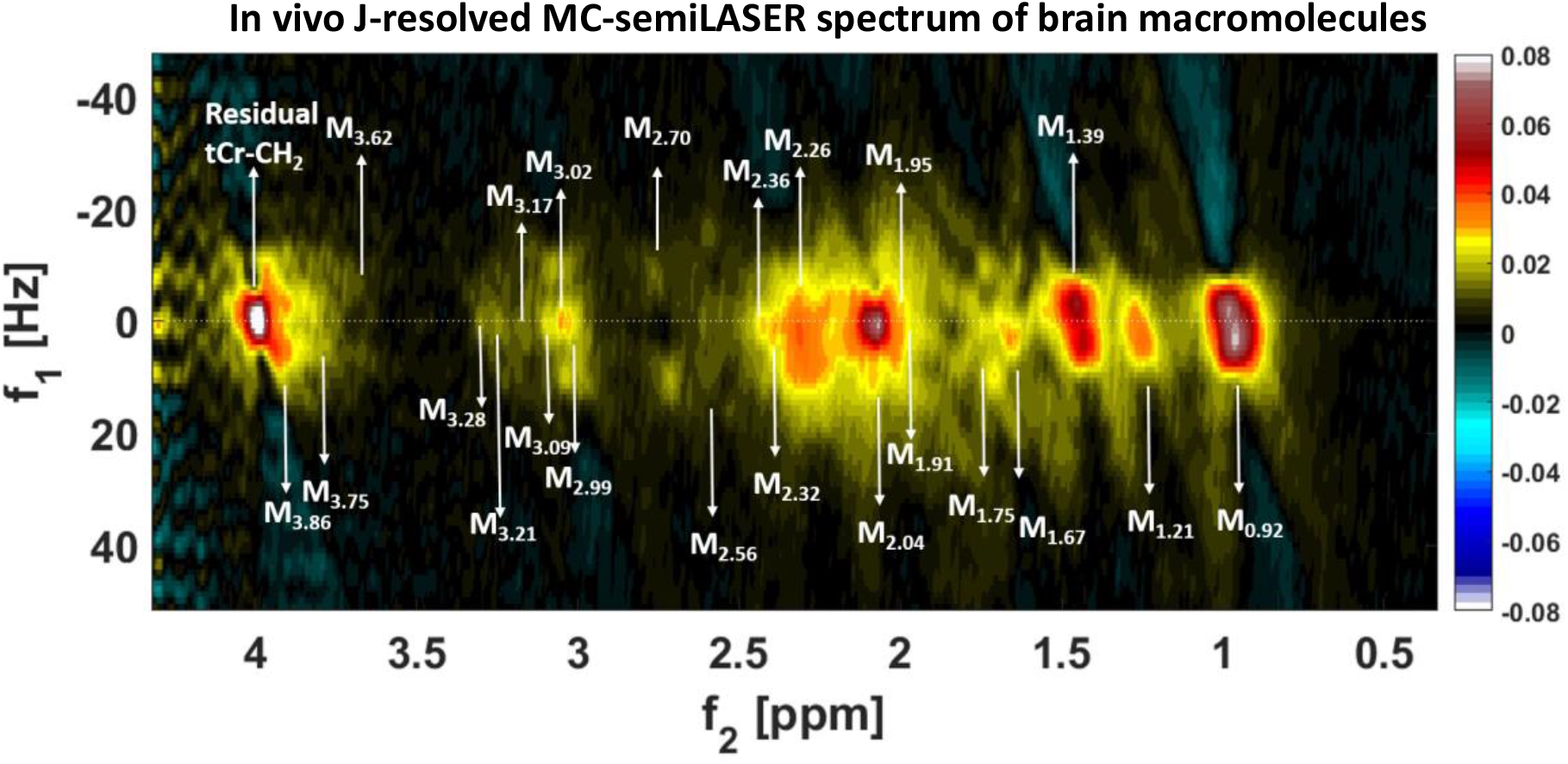
2D DIR J-resolved MC semiLASER (TI_1_/TI_2_: 2360/625 ms, *n* = 60, Δ*t* = 2 ms) summed spectrum from five healthy volunteers. The subscripts in the labelled MM peaks are the chemical shift in ppm at which the respective MM resonance occurs.

### 3.2 Spectral Fitting

A representative metabolite 2D spectral fit is shown in Figure 6. The minimum 2D residual shows good quality fit indicating that the metabolites included in the basis set modeled the acquired data sufficiently well. The fit quality was similar for all the datasets.

**Figure 6.**
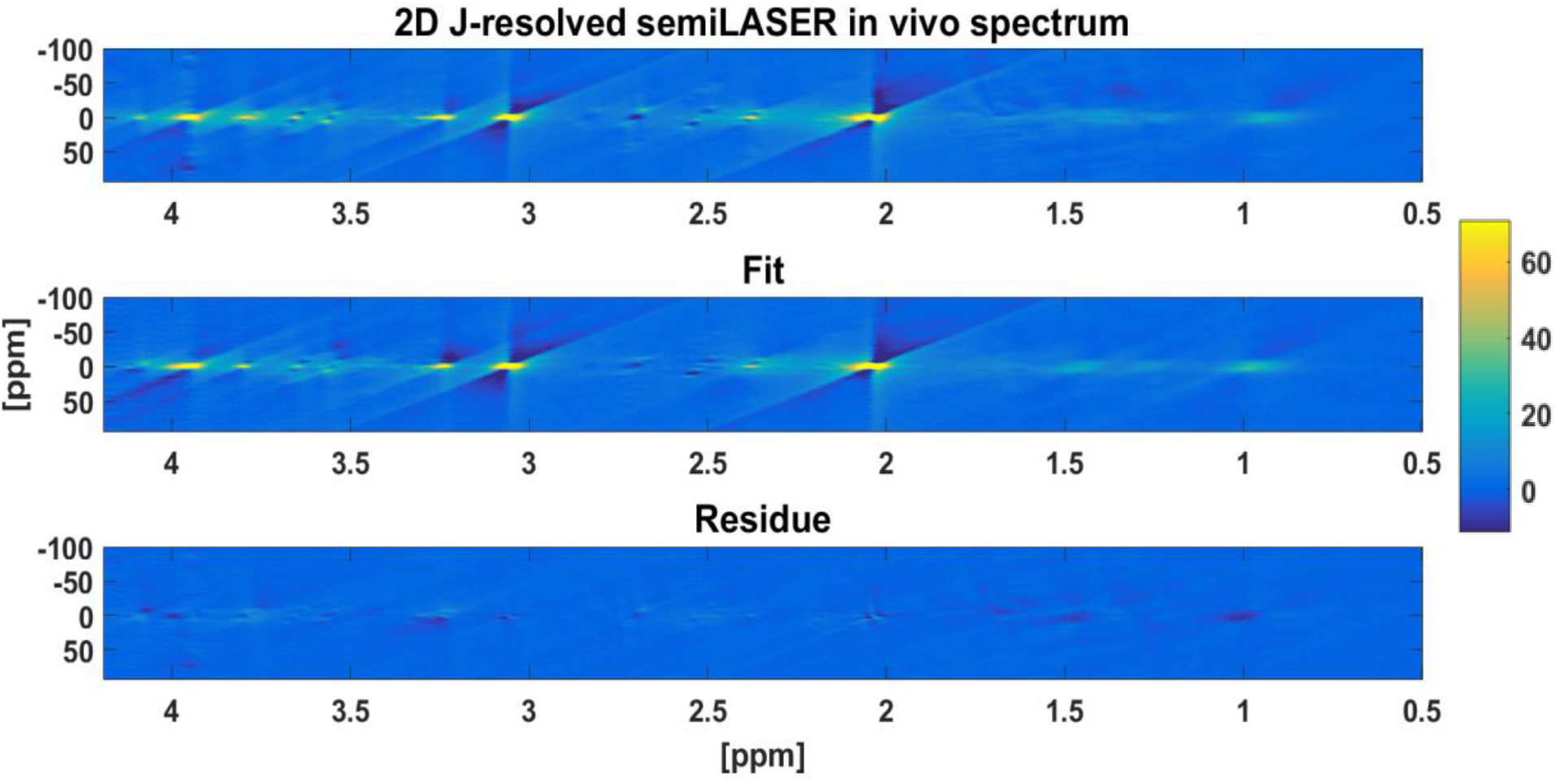
A representative fit of 2D J-resolved MC semiLASER data acquired at 9.4 T. The figure shows the data, the fit and the residual from top to bottom scaled similarly. The fitting was performed in ProFit 2.0 using a tailored 2D basis set created using VesPA.

**Figure 7.**
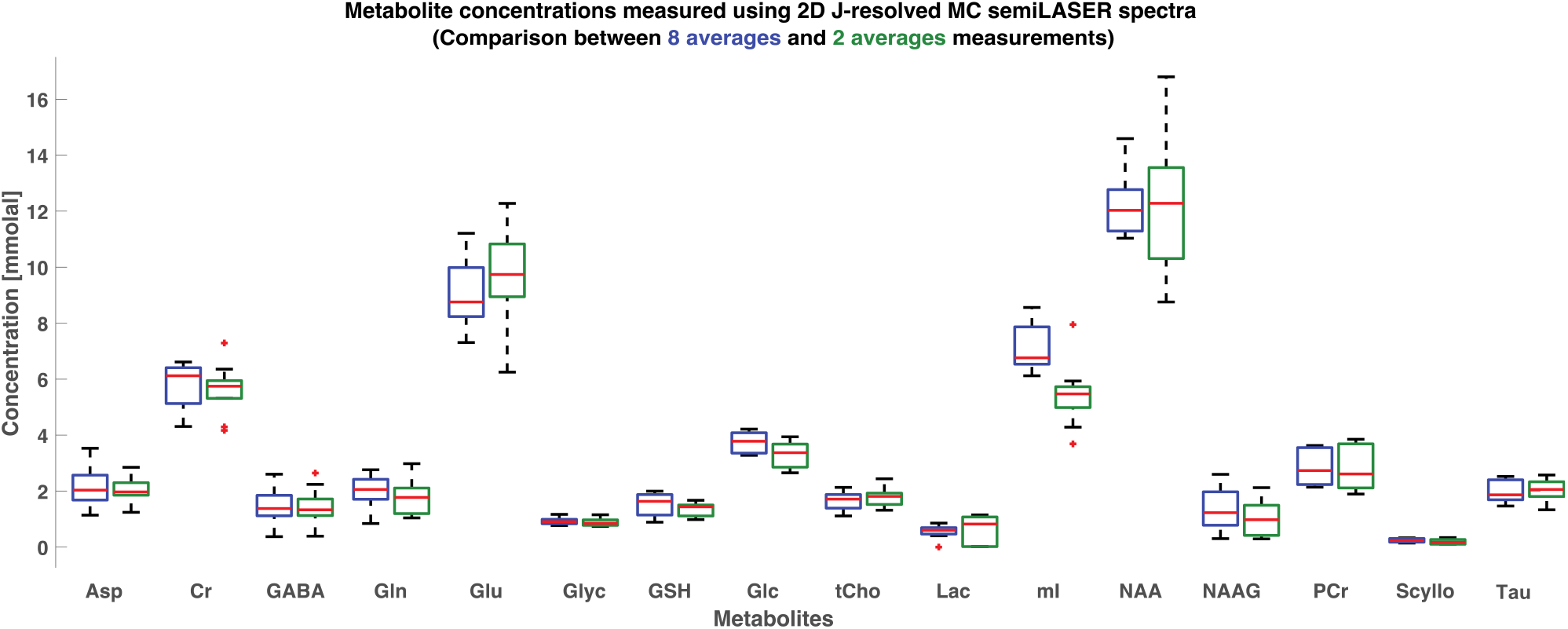
Box plots of metabolite concentrations in mmol/kg measured using 2D J-resolved MC semiLASER spectra (with 8 averages and 2 averages per TE). No significant differences were noted observed in concentrations of metabolites obtained considering 8 or 2 averages per TE. Horizontal lines inside the box plots show median values (50% quartile). The bottom and the top boundaries of the boxes indicate 25% and 75% quartiles respectively. Plus signs (+) show outlier values.

### 3.3 Concentrations

Figure 7 shows box plots of concentrations of metabolites calculated in mmol/kg when considering 8 and 2 averages per TE, respectively. The concentration values of the metabolites are calculated from the 2D MRS data both for 8 averages per TE spectra and 2 averages per TE spectra (Table 1).

**Table 1.** Comparison of concentration levels of metabolites reported in the study to literature at ultra-high field. Quantification results from this study in mmol/kg are reported for 16 metabolites after correcting for tissue content and water and metabolite relaxation times. ProFit2.0 accounts for the T_2_ relaxation effects of the metabolites. T_1_ relaxation times at 9.4 T from Wright et al^49^ were used to account for the T_1_-weighting of metabolites.

| Metabolites | Concentration values [mmol/kg] |  |  |  |  |  |  |
| --- | --- | --- | --- | --- | --- | --- | --- |
|  | 2D MRS<br>(8 averages<br>per TE)<br>(This work) | 2D MRS<br>(2 averages<br>per TE)<br>(This work) | Wright 9.4 T<br>(2021) | Murali-Manohar<br>9.4 T (2020) | Mekle 7T<br>(2009) | Deelchand<br>9.4T (2010) | Marjanska<br>7T (2012) |
| Asp | 2.13 ± 0.64 | 1.96 ± 0.48 | 4.3 ± 0.5 | 4.84 ± 1.15 | 2.9 ± 0.5 | 2.1 ± 0.5 | 2.9 ± 0.8 |
| Cr | 5.77 ± 0.80 | 5.59 ± 0.92 | - | 5.7 ± 0.85 | 5 ± 0.3 | 3.2 ± 0.5 | - |
| GABA | 1.37 ± 0.62 | 1.40 ± 0.66 | 1.2 ± 0.8 | 1.87 ± 0.92 | 1.3 ± 0.15 | 1.3 ± 0.4 | 1.5 ± 0.3 |
| Glc | 1.65 ± 0.33 | 1.75 ± 0.32 | - | - | - | - | - |
| Gln | 1.96 ± 0.59 | 1.74 ± 0.60 | 7.9 ± 1.0 | 7.61 ± 0.91 | 2.2 ± 0.4 | 2.2 ± 0.2 | 1.5 ± 0.5 |
| Glu | 9.11 ± 1.21 | 9.57 ± 1.75 | 12.1 ± 1.3 | 10.9 ± 0.8 | 9.9 ± 0.9 | 9.3 ± 0.9 | 9.6 ± 1.3 |
| Glyc | 0.93 ± 0.11 | 0.86 ± 0.13 | 0.9 ± 0.2 | 1.11 ± 0.28 | - | - | 0.7 ± 0.1 |
| GSH | 1.51 ± 0.39 | 1.34 ± 0.24 | 1.5 ± 0.3 | 1.72 ± 0.16 | 1.3 ± 0.2 | 1.1 ± 0.3 | 1.1 ± 0.1 |
| Lac | 0.55 ± 0.22 | 0.63 ± 0.18 | - | - | 0.7 ± 0.1 | 0.5 ± 0.1 | 0.8 ± 0.2 |
| ml | 7.05 ± 0.79 | 5.42 ± 1.12 | 5.5 ± 0.7 | 5.22 ± 0.45 | 5.7 ± 0.5 | 5.3 ± 0.4 | 6.4 ± 0.8 |
| NAA | 12.20 ± 1.03 | 12.11 ± 2.33 | 11.9 ± 0.8 | 12.61 ± 1.02 | 11.8 ± 0.2 | 13.5 ± 1.6 | 12.6 ± 1.7 |
| NAAG | 1.35 ± 0.73 | 0.99 ± 0.62 | 1.7 ± 0.3 | 1.42 ± 0.19 | 1.1 ± 0.4 | 1.1 ± 0.5 | 0.4 ± 0.2 |
| PCr | 2.86 ± 0.61 | 2.79 ± 0.78 | - | 4.43 ± 0.83 | 2.2 ± 0.19 | 3 ± 0.3 | - |
| Scyllo | 0.24 ± 0.06 | 0.17 ± 0.89 | 0.2 ± 0.1 | 0.13 ± 0.06 | 0.3 ± 0.12 | 0.3 ± 0.2 | 0.4 ± 0.1 |
| Tau | 1.96 ± 0.37 | 2.0 ± 0.41 | 1.7 ± 0.5 | 1.58 ± 0.37 | 1.5 ± 0.3 | 1.3 ± 0.2 | 2.1 ± 0.3 |
| tCho | 3.75 ± 0.38 | 3.35 ± 0.27 | 2.9 ± 0.8 | 3.31 ± 0.95 | 3.6 ± 0.3 | 2.5 ± 0.5 | - |
| tCr | 8.64 ± 1.03 | 9.18 ± 1.44 | 9.5 ± 1.0 | 10.10 ± 0.46 | 8.0 ± 0.4 | 7.7 ± 0.4 | 8.7 ± 1.1 |

No significant differences were observed in concentrations of metabolites obtained considering 8 or 2 averages per TE.

## 4. Discussion

### 4.1 Spectral Quality

The phantom tests performed to optimize the step size (Δ*t*) along with the knowledge of T_2_ relaxation times of metabolites in vivo at 9.4 T from Murali-Manohar et al.,^37^ led to the choice of Δ*t* = 2 ms and number of steps, *n* = 85 for acquisition in vivo. There was sufficient sampling (500 Hz) in the indirect dimension to cover the frequency range of interest in all cases. The final choice of the scan parameters was made since the appearance of ‘t_1_-ridges’^21^ due to t_1_ noise and sinc character from t_1_ truncation was diminished significantly by longer TE_max_ keeping in mind also comfortable scan durations. Furthermore, T_2_-weighting signal loss was avoided largely by setting the first echo time TE_min_ to be 24 ms, as short as possible. According to Murali-Manohar et al^37^, the T_2_ relaxation times in the gray-matter rich voxel present in the occipital lobe ranged from ∼45 to 110 ms. Thus, the chosen parameters swept a decent range of TEs from 24 to 194 ms.

Using the adiabatic J-resolved semiLASER localization sequence proposed herein resulted in a chemical shift displacement of 5% per ppm for each voxel dimension (bandwidth: 8000 Hz). Lin et al^6^ showed that the intensity of the J-refocused peaks can be reduced by a factor of

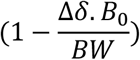

where Δ*δ* is the chemical shift difference in ppm of the spins A and X in an AX spin system, *B*_0_ is the frequency of the static magnetic field in MHz and *BW* is the bandwidth of the refocusing pulse. Considering lactate coupled peaks at 1.31 and 4.09 ppm, the resulting reduction in the intensity of the J-refocused peaks is 86%. Therefore, there were barely any J-refocused peaks in phantom spectra (Figure 3) and no J-refocused peaks in in vivo spectra (Figure 4) that usually appear because of spatially dependent evolution of the J-coupled peaks^21^ due to limited bandwidth refocusing pulses. Consequently, the J-resolved multiplet peaks retained maximum possible intensity.

At 3T typically 8 averages per TE step and 100 TE steps are needed to yield a high quality 2D J-resolved spectrum in about 20 min of measurement time^30^. Hence, 2D J-resolved MC-semiLASER at 9.4T were acquired with 8 averages per TE (acquisition duration of 68 min) and a subset of 2 averages per TE was also investigated (acquisition duration: 17 minutes). It was shown herein that 2 averages per TE (acquisition duration: 17 minutes) yielded the same information as the respective 8 average per TE data set when using metabolite-cycling. Thus, the acquisition time is very similar to 3T 2D J-resolved MRS, which is successfully applied to neuroscientific and clinical studies^8-14^. Considering the high SNR obtained at 9.4T even 1 average per TE would suffice yielding an acquisition duration of only 8 minutes similar to 1D MRS, but in that case a water suppression technique is required instead of employing the 2-step MC method, which requires at least 2 averages per TE. Even though the TR had to be increased at 9.4 T when compared to 3 T due to SAR, the SNR benefit at UHF allows us to reduce the number of averages in turn bringing down the total acquisition comparable to 2D MRS at 3 T^30^ (∼17 minutes) or potentially even lower if no MC was implemented (8 minutes).

Altogether, the application of 2D MRS at UHF is thus feasible for clinical and neuroscientific studies. It is known that the additional benefit at 9.4 T comes from the dispersed chemical shift displacement making the peaks overlap less compared to 3 T. This benefit maximizes when using 2D MRS with MES scheme when the overlap of the J-coupling peak is further reduced. This enabled us to detect peaks such as glucose, lactate and glycine, which are not easily visible in 1D MR spectra even from 9.4 T.

In addition to metabolite spectra, 2D J-resolved DIR MC-semiLASER^38,43^ (acquisition duration: 128 minutes, averages per TE: 16, *n*: 60) sequence was also used to obtain the macromolecular spectra. This MM data was used to account for the underlying MM in the spectral fitting process^47^. MM data was acquired on 5 volunteers.

The knowledge of T_2_ relaxation times of MM (approximately ranging from 15 to 37 ms in GM-rich voxel) at 9.4 T from Murali-Manohar et al^37^ was utilized in setting *n* = 60 (covering TE: 24 to 144 ms) for the acquisition of MM spectra using DIR MC J-resolved semiLASER. By TE_max_ = 144 ms, MM signal decayed completely. Therefore, for *n* = 61 to 85 no corresponding MM spectra were provided in the basis set for metabolite spectral fitting. This study presents a 2D J-resolved MM spectrum (Figure 5) for the first time in vivo in the human brain. M_0.92_, M_1.21_, M_1.39_, M_1.67_, M_1.75_, M_2.04_, M_2.26_, M_2.56_, M_2.70_, M_2.99_ and M_3.86_ are observed to undergo J-evolution and they appear as multiplets in the 2D J-resolved spectrum shown in Figure 5 indicating that these peaks have J-coupled spin systems. This observation agrees with Behar et al^45^ who reported the above-mentioned peaks (except M_2.56_ and M_2.70_) as J-coupled MM observed from COSY and J-resolved spectra of dialyzed human cerebral cytosol at 8.4 T. Giapitzakis et al^38^ reported M_2.56_ and M_2.70_ peaks for the first time and assigned them to β-methylene protons of aspartyl groups which correspond to doublet-of-doublets. This manuscript presents in vivo 2D MM spectra for the first time. Figure 5 also shows M_1.75_, M_1.91_, M_1.95_, M_2.32_, M_2.36_, M_3.02_, M_3.09_, M_3.17_, and M_3.28_ peaks which are observed here in 2D MRS, but not in 1D MRS^37,38,43^ in the human brain at 9.4 T. These MM peaks other than M_1.75_, M_2.32_ and M_3.28_ were previously only reported at 17.2 T in rat brain by Lopez et al^46^.

Thus, 2D MRS at UHF holds the potential to answer some basic research questions such as assigning unlabeled peaks or in understanding the overlap and J-coupling behavior of the MM peaks.

### 4.2 Spectral Fitting

Bandwidth of the refocusing pulse was taken into account while simulating the basis sets in VesPA in order to simulate J-refocused peaks if any, as suggested earlier^21^. The basis sets simulated to fit the 2D metabolite spectra using VesPA also included real pulse shapes with exact durations, bandwidths and the MES scheme. It can be seen from Figure 6 that the simulated 2D basis set fits the data very well.

Spectral fitting of 2D J-resolved data using Profit 2.0 can also be used to determine T_2_ relaxation times of even low-concentration J-coupled metabolites as shown by Wyss et al^48^, which is the subject of a future study.

### 4.3 Metabolite Concentrations

This study quantifies and reports concentration values of 16 metabolites in mmol/kg using an adiabatic 2D J-resolved localization technique at 9.4T in the human brain for the first time. Concentration values of metabolites calculated while considering 8 and 2 averages per TE for the 2D MRS spectra with no significant differences between the values (Figure 7). The calculated millimolal concentration values lie within the range of values that were reported in previous literature^37,41,49-51^ for most of the metabolites. Mekle et al^41^ and Deelchand et al^51^ did not perform T_2_ relaxation correction. However, these studies used shorter TE times; therefore, the contribution from T_2_ weighting may have been insignificant. All the studies included experimentally acquired MM spectrum during the fitting procedure. Mekle et al^41^ set tCr to 8 mmol/kg and used it as internal reference standard.

For metabolites with J-coupled spin systems, low peak amplitudes and/or overlap with larger singlets such as GABA, GSH, NAAG and Tau the concentrations measured from this study are in line with previous 1D MRS studies at 7T^41,50^ and 9.4T^37,49^. Moreover, the 2D MRS technique could quantify Glc and Lac,, which are otherwise challenging to quantify. 2D MRS also allowed for the separation of Glycine and mI. Separation of PCr and Cr was achieved in this study along with two previous 9.4T study^37,49^ and two previous 7T studies. The concentration of Asp measured in this study is in line with two out of three previous 9.4T^49,51^ and two 7T human brain studies^41,50^. Gln concentrations from this study are closely matching those measured in two previous 7T studies^41,50^ and one 9.4T study^51^, but deviate from results of two other previous 1D MRS studies at 9.4T^37,49^. Gln concentration estimate from those 1D MRS studies^37,49^ is higher than expected. The higher estimation of Gln in the 1D MRS study could come from negative LCModel spline baseline near Gln as pointed out in the respective studies. Fitting of 1D MRS data using recently proposed ProFit-1D^52^, which uses more advanced spline baseline modeling, may help overcome this issue,

## 5. Conclusion

2D J-resolved MC semiLASER was successfully implemented for human brain application at 9.4 T. The development of a dedicated analysis pipeline allowed for quantification of the tissue concentrations of 16 human brain metabolites including those with lower SNR and J-coupling induced multiplet spectral pattern such as GABA, Gln, Lac, and Glc. It was also shown that this information can be obtained in a scan time that allows application to neuroscientific and clinical studies in future. In addition, the 2D resolved macromolecular spectrum allows for the observation of additional peaks M_1.75_, M_1.91_, M_1.95_, M_2.32_, M_2.36_, M_3.02_, M_3.09_, M_3.17_, and M_3.28_ peaks, which are not detectable with 1D MRS even at 9.4T.

## Abbreviations

UHF: ultra-high field
NMR: nuclear magnetic resonance
2D: two-dimensional
MRS: magnetic resonance spectroscopy
^1^H: proton
DIR: double inversion recovery
MES: maximum echo sampling
SAR: specific absorption rate
SNR: signal-to-noise ratio
MC: metabolite-cycling
MM: mobile macromolecule

## Acknowledgement

The authors would like to thank Brian Soher for providing the Vespa simulation code for semiLASER sequence. Financial support by the Horizon 2020/ CDS-QUAMRI Grant number: 634541 (A. Henning, T. Borbath, and S. Murali-Manohar), SYNAPLAST Grant number: 679927 (A. Henning, A.M. Wright, and S. Murali-Manohar), and Cancer Prevention and Research Institute of Texas (CPRIT) Grant number: RR180056 (A. Henning) are gratefully acknowledged.

## Supplementary figure captions

**Supplementary figure S1:** (Top to bottom) Phantom spectrum with TE_max_ = 100, 99, and 100 ms for Δ*t*: 2 ms (*n*: 50), 3 ms (*n*: 33), and 4 ms (*n*: 25) respectively. SNR is higher when Δ*t*: 2 ms compared to 3 and 4 ms; t_1_ ridges are not seen to be impacted.

**Supplementary figure S2:** Two-dimensional J-resolved MC semiLASER spectrum in magnitude mode from a representative subject. This spectrum was created considering 2 averages per TE (scan duration: 17 minutes).

