## Supplementary Information for "Quantitative 2D J-resolved metabolite-cycled semiLASER spectroscopy of metabolites and macromolecules in the human brain at 9.4 T"

**Supplementary figure S1:** (Top to bottom) Phantom spectrum with $TE_{max}$ = 100, 99, and 100 ms for $\Delta t$: 2 ms ($n$: 50), 3 ms ($n$: 33), and 4 ms ($n$: 25) respectively. SNR is higher when $\Delta t$: 2 ms compared to 3 and 4 ms; $t_1$ ridges are not seen to be impacted upon changing $\Delta t$ and keeping $TE_{max}$ similar between the measurements.

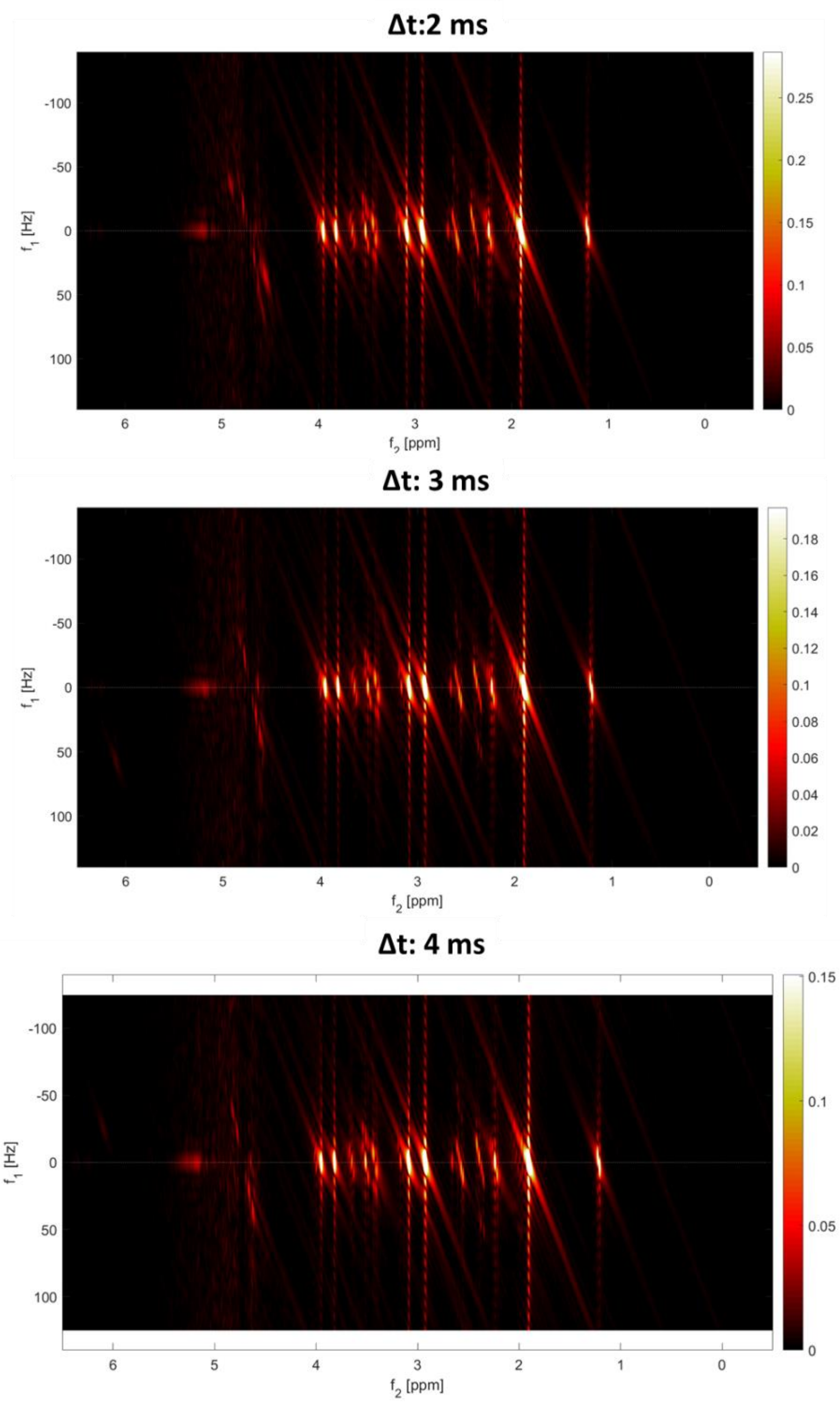

**Supplementary figure S2:** Two-dimensional J-resolved MC semiLASER spectrum in magnitude mode from a representative subject. This spectrum was created considering 2 averages per TE (scan duration: 17 minutes).

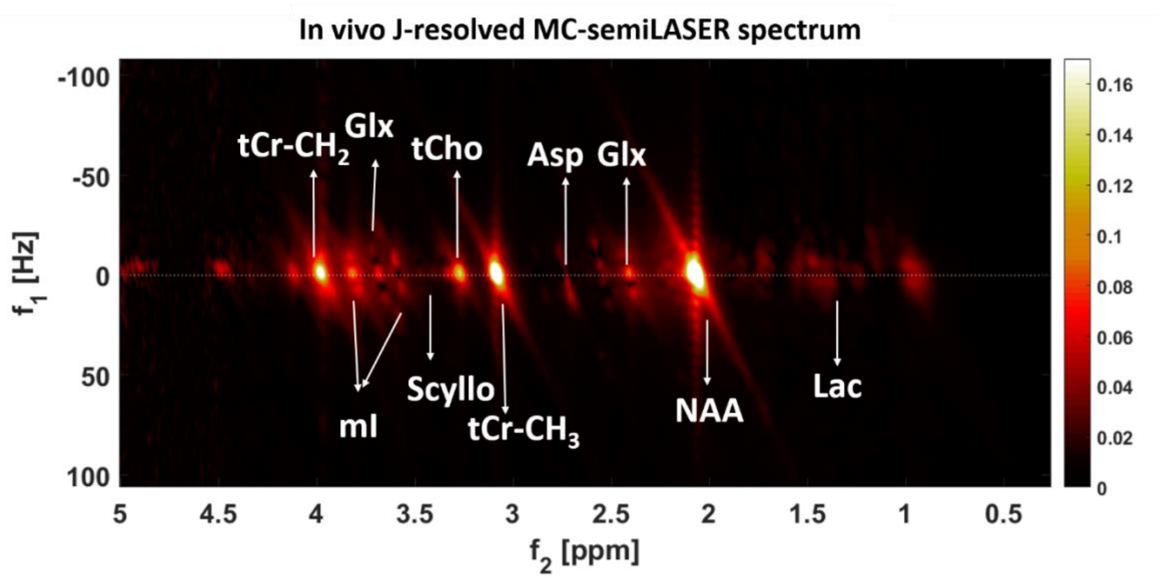


### Annex A – Quantification of metabolites

Metabolite concentrations $[M]$ were quantified in mmol/kg from the ProFit 2.0 concentration results after fitting the metabolite spectra acquired from all subjects.

For internal water referencing, the concentration values from fitting were corrected for tissue water fractions and relaxation times as follows:

$$[M] = \frac{S_{Met} \times (f_{GM} \times R_{H20_{GM}} + f_{WM} \times R_{H20_{WM}} + f_{CSF} \times R_{H20_{CSF}})}{S_{H20}\,(1 - f_{CSF}) \times R_{Met}} \times \frac{2}{(1 + F_S)} \times [H_2O]$$

where $f_y = \frac{f_{y\_vol} \times a_y}{f_{GM\_vol} \times a_{GM} + f_{WM\_vol} \times a_{WM} + f_{CSF\_vol} \times a_{CSF}}$

Here $y$ corresponds to either $GM, WM$, or $CSF$; $f_{y\_vol}$ is the fraction of the respective tissue type determined by segmentation; $a_y$ are the relative densities of MR-visible water for the given tissue types (78%, 65%, 97% for $GM, WM$ and $CSF$ respectively); these $a_y$ values were further scaled for the relative densities of GM and WM tissue (1.04 g/ml)[1–4]. To arrive at mmol/kg units , the concentration of the MR-visible water [$H_2O$] within a voxel was considered in molal concentration and  assumed to be that of pure water (55,510 mmol / kg)[5]. $S_{Met}$ is the signal from the metabolite peak.

ProFit 2.0 accounts for the $T_2$ relaxation times of water and metabolites from the second TE to the last TE by including respective line shape models specific to each metabolite in the fitting algorithm. Therefore, $T_2$ correction was included only for the first TE = 24 ms.

$$R_{H2O\_y} = \left[1 - \exp\left[-\frac{TR}{T1_{H2O_y}}\right]\right] \exp\left[-\frac{TE}{T2_{H2O_y}}\right]$$

is the relaxation correction factor for each tissue type $y$. $T1_{H2O_y}$ is the $T_1$ relaxation time of water in the tissue type $y$; in particular, the $T_1$ relaxation times of water in GM are $T1_{H20\_GM}$ = 2120 ms; in WM are $T1_{H20\_WM}$= 1400 ms; and in CSF are $T1_{H20\_CSF}$ = 4800 ms at 9.4 T[6]. $T2_{H2O_y}$ is the $T_2$ relaxation time of water in the tissue type $y$; $T2_{H2O\_GM}$ = 37 ms, $T2_{H2O\_WM}$ = 30 ms, and $T2_{H2O\_CSF}$ = 181 ms.

$$R_{Met} = \left(1 - \exp\left[-\frac{TR}{T1_{Met}}\right]\right) \left[-\frac{TE}{T2_{Met}}\right]$$

is the relaxation correction term for metabolites. $T1_{Met}$ values from were taken Wright et al[7] and $T2_{Met}$ values were taken from Murali-Manohar et al[8]. The denominator $1 - f_{CSF}$ was implemented for partial-volume correction arising from contributions of CSF to the voxel volume. The factor $\frac{2}{1+Fs}$ was introduced to correct for the multiplication of even numbered acquisitions with the scaling factor ($F_s$) from metabolite cycling.
